# Using supervised machine learning to quantify cleaning behaviour

**DOI:** 10.1101/2025.06.03.657613

**Authors:** Raul Oliveira, Nuno Cruz Garcia, José Ricardo Paula

## Abstract

Cleaner fish engage in mutualistic interactions by removing ectoparasites from client species, a behaviour that has traditionally been quantified through labour-intensive manual video analysis. This method is not only time-consuming but also susceptible to human error and bias. In this study, we developed a semi-automated system to track and classify cleaning interactions between the cleaner wrasse (*Labroides dimidiatus*) and the powder blue tang (*Acanthurus leucosternon*) in a controlled three-dimensional laboratory setting. We employed DeepLabCut (DLC), a deep learning-based tool for markerless pose estimation, to track both fish species simultaneously. The resulting model reliably tracked both individuals with low error rates. Using the tracking data, we designed a classification algorithm that detected cleaning interactions with 90% accuracy. Although the algorithm misclassified approximately 15% of non-interactions as interactions, it successfully identified 25% of video content as containing interactions, thereby reducing the amount of footage requiring manual annotation by 75%. This approach significantly decreases human labour while maintaining high classification performance. Overall, our system represents a valuable step toward automating behavioural analysis in marine mutualisms and can serve as a foundation for broader applications in ethology and conservation research.

## 1. Introduction

In coral reefs, cleaner wrasses are well-known for their cleaning activity, where they inspect client fish and feed on ectoparasites removed from their (Grutter, 2002). This mutualistic interaction is critical to the health of reef ecosystems, with cleaner fish interacting with more than 2,000 clients daily, and some clients seeking out cleaners for interaction up to 145 times a day (Grutter, 1996). Cleaning interactions include a a wide array of behaviours such as advertising dances, manipulation, tactile stimulation, cleaning bites, client jolts, chasing, punishment – all of which provide insight into cleaning motivation and interaction quality (Paula et al., 2019). Among cleaner fish, *Labroides dimidiatus*, the Indo-Pacific cleaner wrasse, stands out as one of the most extensively studied species. It has become a model system for investigating the evolution of interspecies cooperation and has contributed significantly to our understanding of coral reef dynamics (Demairé et al., 2020; Soares, 2017).

Traditionally, interspecific interactions are analysed through manual video recording processing (Lindburg, 1969; Pollok et al., 2000; Reiss & Marino, 2001). While widely used, this method is labour-intensive and often unreliable due to variability among observers and the effects of fatigue from prolonged video analysis (Baker, 2016). To mitigate these issues, some journals now require interobserver reliability assessments to validate behavioural data (Anderson & Perona, 2014; Arac et al., 2019; Dell et al., 2014; Gomez-Marin et al., 2014). However, due to fluctuating attention levels, even trained observers may miss or misclassify behaviours, leading to missed detections (Kays et al., 2015). In addition to these analytical challenges, behaviour itself is inherently complex, dynamic, and multidimensional, often exceeding the descriptive power of traditional methods (Anderson & Perona, 2014). These limitations underscore the need for innovation in behavioural research methodologies. Recent advances in computer vision and artificial intelligence have significantly enhanced the field, enabling more precise, efficient, and scalable analyses of animal behaviour (M. W. Mathis & Mathis, 2020). By automating tracking and reducing dependence on human observation, these tools minimize bias, reduce labour, and improve the overall accuracy of behavioural datasets (Berman, 2018; Dell et al., 2014).

Since the early 1990s, research projects focused on motion tracking, facial pattern recognition, and human detection (Yang & Huang, 1994). In the area of computer vision, pose estimation (PE) refers to the task of localizing the joint regions or set of keypoints of an object (e.g., fish, car, human) within an image (Cao et al., 2019). The use of PE has steadily expanded into everyday applications, including gesture-based human–computer interaction (Nguyen et al., 2020), assessment and correction of human movement and posture in healthcare and sport applications (Chen & Yang, 2020), among others. The emergence of deep learning has catalyzed rapid advancements in pose estimation, opening new domains such as animal-specific PE (A. Mathis et al., 2018). Animal pose estimation is increasingly important for understanding behavioural dynamics (Joska et al., 2021), monitoring wildlife migration (Bauer & Klaassen, 2013), and even in pet care applications like automated dog monitoring (Biggs et al., 2020). While pose estimation models for humans and animals often share a common underlying architecture (Graving et al., 2019), the high biological diversity across species and the scarcity of labeled datasets pose unique challenges for the development of robust animal PE models (Pereira et al., 2019).

As automatic posture tracking and behavioural classification become increasingly central to the study of animal behaviour across species (e.g., humans, mice, and pigeons;) (Alghamdi et al., 2015; Arac et al., 2019; Wittek et al., 2022; Zhang et al., 2020), there is a growing demand for open-source tools capable of accurately measure and classify behavioural interactions. To date, most pose-estimation research has focused on single animals in two-dimensional environments, typically involving model species such as mice, flies, or zebrafish (Bohnslav et al., 2021; Gerós et al., 2020; Guilbeault et al., 2021; Jia et al., 2022). Studies that explore behaviour in three-dimensional spaces usually do so with individuals from the same species (Arablouei et al., 2023; Han et al., 2024; Long et al., 2020). However, to the best of our knowledge, no existing tools are capable of reliably track and classify interspecific cleaning interactions between the cleaner wrasses and their client fish. In this study, we used the machine-learning-based tracking software DeepLabCut (DLC) as a pose-estimation tool (A. Mathis et al., 2018; Nath et al., 2019) to simultaneously track two distinct species—the cleaner wrasse (*Labroides dimidiatus*) and its client fish (*Acanthurus leucosternon*)—by assigning species-specific keypoints. Building upon this tracking data, we developed a classification algorithm designed to automatically detect cleaning interactions based on spatial proximity, orientation, and characteristic movement patterns associated with cleaning interactions.

In summary, this study presents three main contributions: (i) the first multi-species 3D DeepLabCut model for cleaner–client fish; (ii) a novel distance-based classifier with quantitative performance evaluation; and (iii) a 75% reduction in manual effort for video annotation, enabling more scalable behavioural analysis.

## 2. Materials and Methods

### 2.1. Video Acquisition and Video Analysis

#### 2.1.1. Hardware tools for video acquisition

We utilized video recordings from a previous behavioural study conducted in 2018 (Ramírez-Calero et al., 2023). In brief, the experiment involved 24 pairs, each consisting of one cleaner wrasse (*Labroides dimidiatus*) and one client fish (*Acanthurus leucosternon*), placed together in observation tanks (40 L) within a designated observation room. Prior to the behavioural trials, both cleaners and clients underwent a 24-hour fasting period to standardize motivation for interaction. Each trial consisted of introducing a cleaner-client pair into a tank and recording their behaviour for 40 minutes, excluding the initial 5 minutes allocated for acclimation. The video setup included three cameras covering every two aquariums to enable three-dimensional tracking of fish movements. Specifically, one camera was positioned in front of each tank to record lateral views, while a top-mounted camera captured a dorsal view of both aquariums simultaneously (Figure 1). All videos were recorded in full HD resolution (1920×1080 pixels) at a frame rate of 50 frames per second, ensuring high temporal and spatial fidelity for subsequent tracking and analysis.

**Figure 1.**
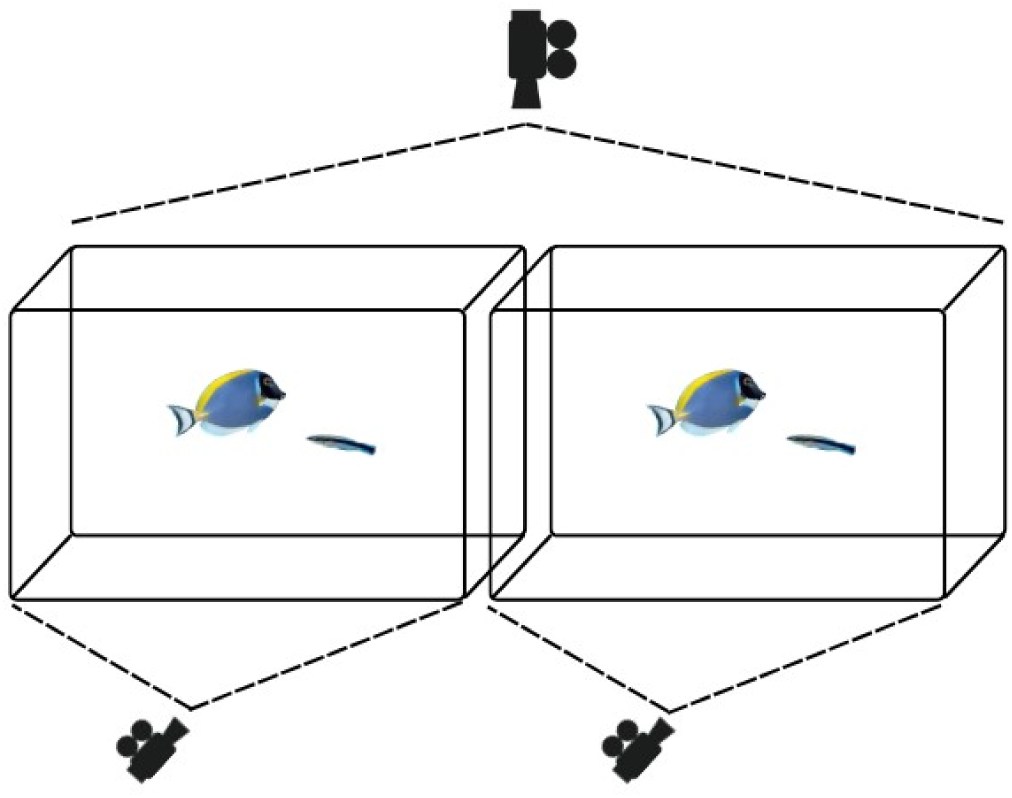
Experimental setup for recording cleaner-client fish interactions. Each behavioural trial was conducted in a pair of adjacent 40 L aquariums, each containing one cleaner wrasse (*Labroides dimidiatus*) and one client fish (*Acanthurus leucosternon*). The interaction between the fish was recorded using three synchronized cameras. One top-mounted camera recorded the dorsal view of both tanks simultaneously, while two side-mounted cameras (one per aquarium) captured the lateral perspective. This multi-angle configuration enabled the reconstruction of fish movements in three dimensions for subsequent tracking and analysis.

#### 2.1.2. Hardware tools for video analysis

All video analyses were performed on a workstation (ThinkStation P620), with a Processor AMD Ryzen™ Threadripper™ PRO 3945WX @ 4,0 GHz - 4,3 GHz, 32GB of RAM, NVIDIA RTX A4000 16GB RAM, running on a Microsoft Windows 11 Pro.

### 2.2. Labelling events

To implement a supervised machine learning approach for both the training and testing stages, interaction events needed to be manually annotated. We used the open-source software BORIS (Behavioural Observation Research Interactive Software; (Friard & Gamba, 2016) to label interaction events.

### 2.3. Pre-Processing Video Data

For the tracking tool to read the video recordings, were re-encoded the original codec of the FrontView movies. To re-encode, we used FFmpeg (FFmpeg Developers, 2016) with the following command line:

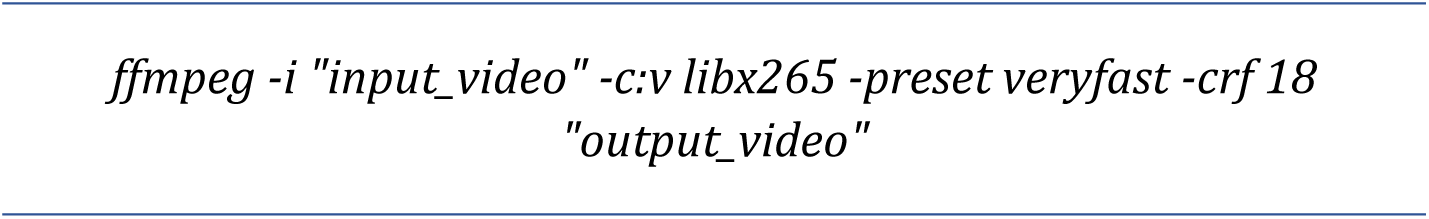

We used the Capcut, a free video editing program, (*CapCut | All-in-One Video Editor & Graphic Design Tool Driven by AI*, n.d.) to split the TopView videos into two parts, one for the left aquarium and the other for the right aquarium.

### 2.4. Tracking tool

To develop the automated tracking system, we used DeepLabCut (DLC) with the dlcrnet_ms5 architecture, employing the default neural network parameters as specified in the official documentation (M. Mathis, 2021). To simultaneously track two distinct species, *Labroides dimidiatus* (cleaner fish) and *Acanthurus leucosternon* (client fish), we assigned unique keypoint groups: the “uniquebodyparts” group was used for cleaner fish tracking, while the “multianimalparts” group was used for the client fish. Each video frame was initially stored in raw (decompressed) format at 1920 × 1080 pixels with 8-bit depth (∼16.5 Mbit per frame). During DLC processing, this input was transformed into a compact format comprising 14 body parts × 2 coordinates × 32-bit precision (896 bits), yielding an approximate 18,500-fold reduction in data bandwidth per frame. To reconstruct three-dimensional trajectories, we trained two independent 2D tracking models corresponding to two different camera perspectives. These were later integrated to enable 3D pose estimation of fish interactions within the observation tanks.

#### 2.4.1. Frame labelling of the FrontView tracking model

To identify significant elements in an image (i.e., recognition of the fish body parts), we used a frame labelling technique. We labelled ten distinct key points on the client fish: Client_Mouth, Client_SpineHead, Client_SpineMid, Client_BodyTop1, Client_BodyTop2, Client_BodyBot1, Client_BodyBot2, Client_Tail, Client_TailTop, and *Client_TailBot* (Figure 2.A). The cleaner fish was labelled with four distinct keypoints: *Cleaner_Mouth*, *Cleaner_Spine1*, *Cleaner_Spine2* and *Cleaner_Tail* (Figure 2.B).

**Figure 2:**
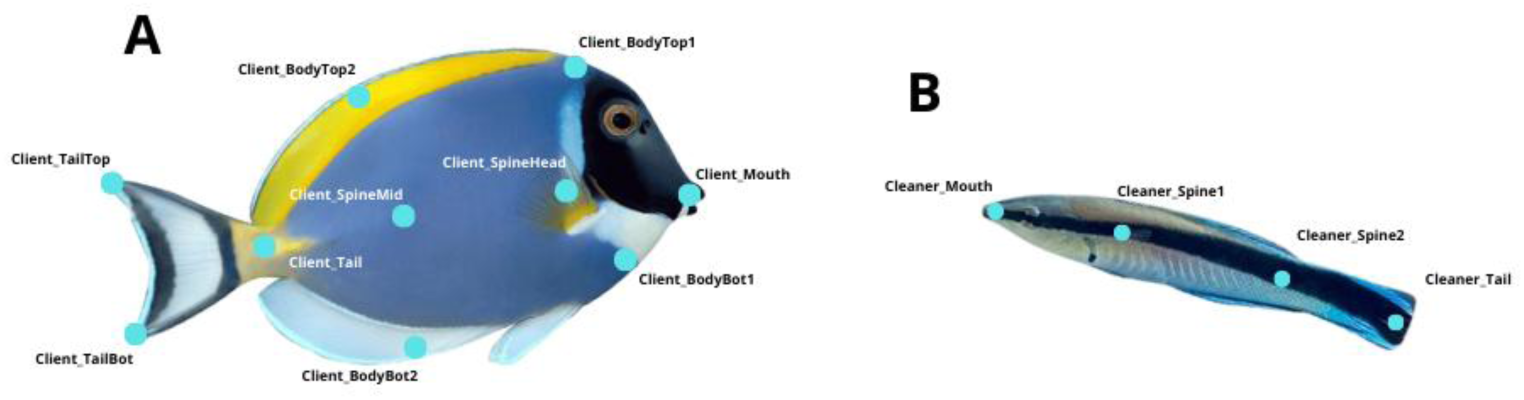
Marked keypoints from the FrontView camera for (A) the client fish, *Acanthurus leucosternon* (n=10), and (B), the cleaner fish, *Labroides dimidiatus* (n=4). These key points are marked whenever visible in the frame.

We manually labelled 20 frames from each of the 20 videos of the FrontView tracking model, for a total of 400 labelled frames. 95% of these frames were used for training the automated model. For this, we used a neural network based on dlcrnet_ms5, as recommended by the documentation for multi-animal models. We traubed the network for 75000 iterations with a batch size of 16 (Figure S1).

#### 2.4.2. Frame labelling of the TopView tracking model

From the top view cameras, we labelled 11 distinct key points for the client fish: *Client_Mouth, Client_EyeL*, *Client_EyeR*, *Client_FinBaseL*, *Client_FinTipL*, *Client_FinBaseR*, *Client_FinTipR*, *Client_Spine1*, *Client_Spine2*, *Client_Tail* and *Client_TailTip* (Figure 3.A). We labelled four distinct keypoints for the cleaner fish: Cleaner_Mouth, Cleaner_Spine1, Cleaner_Spine2, and Cleaner_Tail (Figure 3.B).

**Figure 3.**
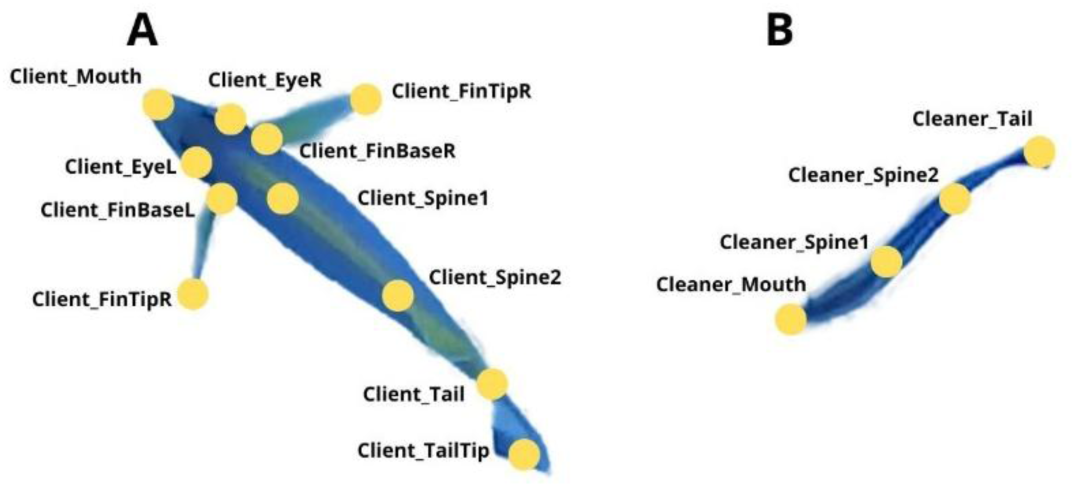
Annotated keypoints from the top-view camera for (A) the client fish, *Acanthurus leucosternon* (n=11), and (B), the cleaner fish, *Labroides dimidiatus* (n=4). These key points are marked whenever visible in the frame.

We labelled 120 frames from each of the six videos for the TopView tracking model, resulting in a total of 720 labelled frames. We used the same parameter as in the Front View model, using 20000 trained iterations (Figure S2).

#### 2.4.3. Missing Values

To address the potential issue of occluded or undetected body parts during tracking, we applied the K-Nearest Neighbors Imputer (KNN Imputer) from the scikit-learn machine learning library (Pedregosa et al., 2011) to impute missing values. This method was selected for its ability to preserve as much of the dataset as possible while maintaining the temporal and spatial coherence of the fish trajectories. Given the dynamic and complex nature of fish movement, KNN imputation helps retain the continuity of behavioural patterns and ensures minimal disruption to downstream classification tasks.

#### 2.4.4. Validation

To ensure the accuracy of our tracking results, we evaluated model performance using three complementary approaches. First, we conducted a qualitative assessment by visually inspecting a subset of labeled videos generated after model training. This manual verification confirmed that the predicted keypoints corresponded accurately to the anatomical landmarks on both fish species, providing a preliminary gauge of model precision. Second, we performed a quantitative evaluation by analyzing the likelihood scores output by DeepLabCut. These scores are derived from the model’s confidence heatmaps and indicate the probability that a predicted keypoint is correctly positioned. By examining these likelihood values across frames, we assessed the overall reliability and consistency of the model’s predictions.

i. **Heatmap Generation**: For each keypoint, the network outputs a heatmap *H*(*x*,*y*), where (*x*,*y*) is the pixel position, and *H*(*x*,*y*) represents the confidence value at that pixel.
ii. **Softmax Normalization**: The raw heatmap values are normalized using the softmax function to ensure they represent probabilities. The normalized heatmap *P*(*x*,*y*) is given by:

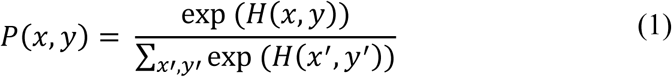

Where exp is the exponential function, and the denomination sums over all pixel positions (x’, y’) in the heatmap.

iii. **Likelihood Calculation**: The likelihood of a keypoint is the value of the normalized heatmap *P*(*x*,*y*) at the predicted keypoint location (*x*_max_,*y*_max_) ), which corresponds to the pixel with the highest intensity in the raw heatmap:

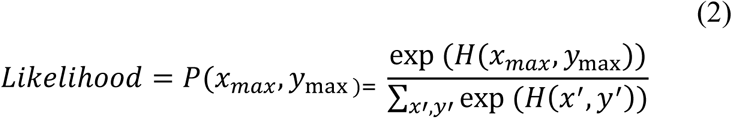

To evaluate the success of model training, we used the likelihood scores provided by DeepLabCut for each labeled body part across all frames. By calculating the average likelihood for each feature, we determined the quality of training, with successful models showing values above 0.90 for every tracked keypoint. Following this, we assessed the quality of model predictions during video analysis by computing the average global likelihood, or model confidence, for each video. This metric is derived by averaging the likelihoods across all features and frames. Videos with model confidence values above 0.95 were considered to be reliably predicted, whereas those with lower values (<0.95) were flagged for potential tracking inconsistencies.

Manual label validation is the third and most reliable method for evaluating model performance. This process involves calculating the Mean Average Error (MAE) between the manually labeled ground truth keypoints (x_i_^true^,y_i_^true^) and the model-predicted keypoints (x_i_^pred^,y_i_^pred^). The MAE is defined as:

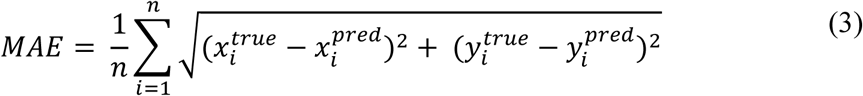

DeepLabCut integrates both missing values (allowing for the exclusion of occluded body parts) and validation approach. The computation output includes: i) all labeled keypoints, and ii) labelled keypoints with a likelihood value greater than a specified threshold p-cutoff (i.e., 0.75 in this study).

#### 2.4.5. Video testing of the newly built automated tool

We tested the newly built automated tool by using six videos from the front view and three from the top view, representing six different interactions (Table S1).

### 2.5. Behavioural classification

Before implementing behavioural classification, we developed a custom function to identify the start and end points of potential interactions. The primary criterion for detecting these events was the proximity between the cleaner and client fish, as mutualistic cleaning interactions can only occur when the individuals are within close spatial range. Given the richness of our dataset, comprising numerous keypoints per individual, we evaluated several strategies to determine the most effective approach for estimating inter-fish distance. These strategies were designed to optimize the prediction of candidate interaction intervals, which were then passed to the classifier for final behavioural identification.

#### 2.5.1.1. Strategy 1 - “C4Cleaner-C4Client”

In strategy 1, we calculated the centroid of the cleaner fish by using all four keypoints (Cleaner_Mouth, Cleaner_Spine1, Cleaner_Spine2 and Cleaner_Tail), and the centroid of the client fish by using four keypoints (Client_Mouth, Client_Spine1, Client_Spine2, Client_Tail). This approach allows the use of other tracking software in the future that utilizes the centroid of the tracking object. Then we calculated the Euclidean distance between the centroids.

#### 2.5.1.2. Strategy 2 - “C4Cleaner-E5Client”

For strategy 2, we calculated the centroid of the cleaner fish using all four keypoints (*Cleaner_Mouth, Cleaner_Spine1, Cleaner_Spine2* and *Cleaner_Tail*). Next, we calculated the Euclidean distance to five keypoints from the client fish (*Client_Mouth, Client_Spine1, Client_Spine2, Client_Tail* and *Client_TailTop)* using the cleaner’s centroid and saved the shortest distance.

#### 2.5.1.3. Strategy 3 - “C2Cleaner-E5Client”

In Strategy 3, we followed the same approach as in Strategy 2 but reduced the number of key points used in the case of the cleaner. We chose to use only two primary points, namely “Cleaner_Mouth” and “Cleaner_Spine1”, as they hold more biological significance (the mouth is the primary organ used by the cleaner for its cleaning activity, and the pectoral fins are used by the cleaner to provide tactile stimulation). We calculated the centroid of the cleaner fish using these two keypoints (*Cleaner_Mouth and Cleaner_Spine1*). Next, we calculated the Euclidean distance to five keypoints from the client fish (*Client_Mouth, Client_Spine1, Client_Spine2, Client_Tail,* and *Client_TailTop)* using the cleaner’s centroid and saved the shortest distance.

#### 2.5.1.4. Strategy 4 - “C2Cleaner-E10Client”

For Strategy 4, we followed the same approach as Strategy 3 but used all ten keypoints from the client fish. Firstly, we calculated the centroid of the cleaner fish based on *Cleaner_Mouth* and *Cleaner_Spine1* keypoints. Then, we measured the Euclidean distance from the cleaner’s centroid to the 10 keypoints of the client fish (i.e., *Client_Mouth*, *Client_Spine1*, *Client_Spine2*, *Client_SpineTop1*, *Client_SpineTop2*, *Client_SpineBot1*, *Client_SpineBot2*, *Client_Tail*, *Client_TailTop*, and *Client_TailBot*) and saved the shortest distance.

#### 2.5.2. Classification equation

To detect cleaning interactions from the tracking data, we developed an equation that analyses a time series of inter-fish distances. The algorithm identifies the onset of a potential interaction when the distance between fish falls below a predefined distance threshold. It then continues tracking subsequent frames to determine whether the interaction persists for a sufficient duration, defined by a minimum number of consecutive frames. If this duration criterion is met, the event is registered as a valid interaction, and the algorithm logs its start and end frames. Additionally, the function checks for temporal proximity between interactions; if two events occur within a defined margin, they are merged into a single interaction instance to account for brief interruptions. At the end of each distance sequence, the algorithm also evaluates whether an ongoing interaction meets the minimum frame requirement before concluding and logging it. The output is a list of tuples, each representing a detected interaction period with the corresponding start and end frame indices. This classification equation relies on two primary parameters, maximum distance and minimum consecutive frames, which allow for robust and efficient identification of cleaning interactions across the dataset.

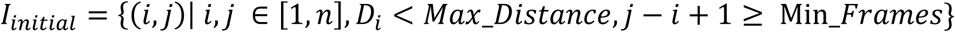

Where:

- *I_initial_* represents the initial set of detected interactions
- (*i*,*j*) denotes an interaction starting at frame *i* and ending at frame *j*
- *D_i_* represents the distance at the *i* -th element of the distance sequence
- The condition *D_i_* < *Max_Distance* ensures that the distance at frame *i* is below the specified threshold
- The condition (*j − i +1*) ≥ *Min_Frames* verifies if the number of consecutive frames from *i* to *j* is greater than or equal to the minimum consecutive frames required for an interaction

To finalize our equation, we performed a merging condition:

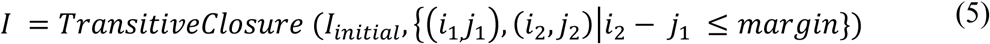

Where:

- *I* represent the final set of detected interactions
- **Merging Condition**: For any two pairs (𝑖_1,_𝑗_1_), (𝑖_2_, 𝑗_2_) ∈ *I_initial,_*, if 𝑖_2_ − 𝑗_1_ ≤ 𝑚𝑎𝑟𝑔𝑖𝑛 and 𝑖_1_ < 𝑖_2_, merge them into a single pair (𝑖_1,_𝑗_2_)
- **Update I:** Replace (𝑖_1,_𝑗_1_) and (𝑖_2,_𝑗_2_) with (𝑖_1,_𝑗_2_) in *I* and repeat until no more pairs satisfy the merging condition.

##### 2.5.2.1. Parameters used for behavioural classification

We conducted an exploratory analysis to determine the distance threshold and interaction length before deciding on the function’s parameters. We observed that there is an increased number of interactions with a duration of interaction between the fish of less than 2 seconds (Figure S3, Table S2).

Since the classification equation depends on the minimal duration while the fish are near one another, we set the *Min_Frames* parameter to 30, 45, or 60, which corresponds to 1, 1.5, and 2 seconds, respectively.

We performed a comparative distribution of the distances for each strategy regarding the *Max_Distance* parameter (Figure S4). We used distinct sets of distances for each strategy since they relied on different keypoints for measuring the distances. For strategy 1 and 2 we used 150, 175, 200, 225 and 250 pixels of distance, whereas we used 100, 125, 150, 175, 200, 225 and 250 pixels of distance for strategy 3 and 4.

For the *margin* parameter, we chose either: i) *margin* = 30, if there is a gap of less than 30 frames (1 second) between the predicted interactions, they will be combined into a single interaction, or ii) *margin* = 0, this rule does not apply.

##### 2.5.2.2. Equation Performance

To evaluate the performance of the behavioural classification function, we employed a set of standard classification metrics: accuracy, precision, recall, specificity, and F1-score. These metrics provided a comprehensive assessment of the algorithm’s ability to identify true interactions while minimizing false detections correctly. In addition to these quantitative indicators, we conducted a qualitative inspection of the classification outputs. Specifically, we examined the total number of correctly and incorrectly predicted interactions, as well as the frequency of duplicate predictions, where a single interaction was erroneously classified as multiple separate events. These duplicates were used to assess the algorithm’s temporal resolution and segmentation reliability (see Figure S5).

To determine potential differences in performance metrics, we performed a comparative analysis for each strategy between the parameters *margin* = 0 and *margin* = 30. To analyse the data, four statistical tests were employed (i.e., Pearson’s Chi-Squared Test, Adjusted Pearson’s Chi-Squared Test, Two Independent Sample t-Test with Pooled Variance, Z-Test for Differences in Proportions), chosen to address the challenges posed by the small population sizes in this study. These tests were selected following guidelines from prior research on statistical methods appropriate for limited datasets (D’Agostino et al., 1988; Upton, 1982). Therefore, we compared the proportions of the Correct Predictions and the Incorrect Predictions (Table S3).

1. **Pearson’s Chi-Squared Test**: To evaluate the independence of categorical variables, providing a baseline measure of association between key variables in the dataset.

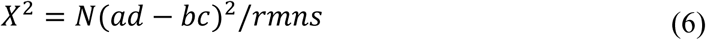

2. **Adjusted Pearson’s Chi-Squared Test**: Due to the small sample sizes, an adjusted version of the Pearson test was employed to account for potential inaccuracies in the Chi-Squared approximation. This adjustment improved the reliability of the p-values for assessing statistical significance.

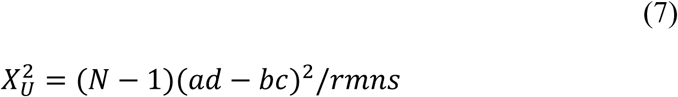

3. **Two Independent Sample t-Test with Pooled Variance**: To compare the means of two groups under the assumption of equal variances. The pooled variance approach was chosen to enhance statistical power despite the constraints of small sample sizes.

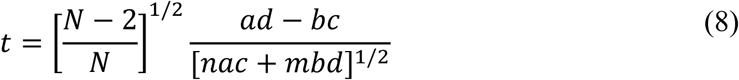

4. **Z-Test for Differences in Proportions**: To analyze the proportion of predicted correct outcomes relative to the total corrected outcomes across groups. The z-test was selected for its efficacy in evaluating proportional differences even under limited population conditions.

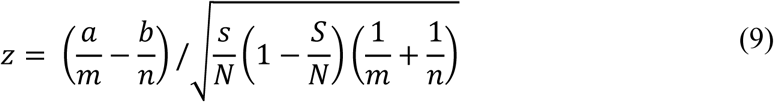

These tests allowed to identify any notable differences in the efficacy and performance of the constructed model by comparing the proportion of correct predictions under the two *margin* settings. This method helped to clarify how changes to the *margin* parameter affected the classification results, which helped to guide improvements of the model.

## 3. Results

### 3.1. DeepLabCut results

#### 3.1.1. Results of FrontView setup

The model created for the FrontView camera presented a low test error after training (Table S4). The average likelihood of all body parts reached 0.92. The Root Mean Square Error (RMSE) values for each body part (labeled by DeepLabCut) were visualized using violin plots and are presented in Figure 4 to compare tracking accuracy. The low RMSE values and high likelihood indicates that DLC managed to accurately track both fish.

**Figure 4.**
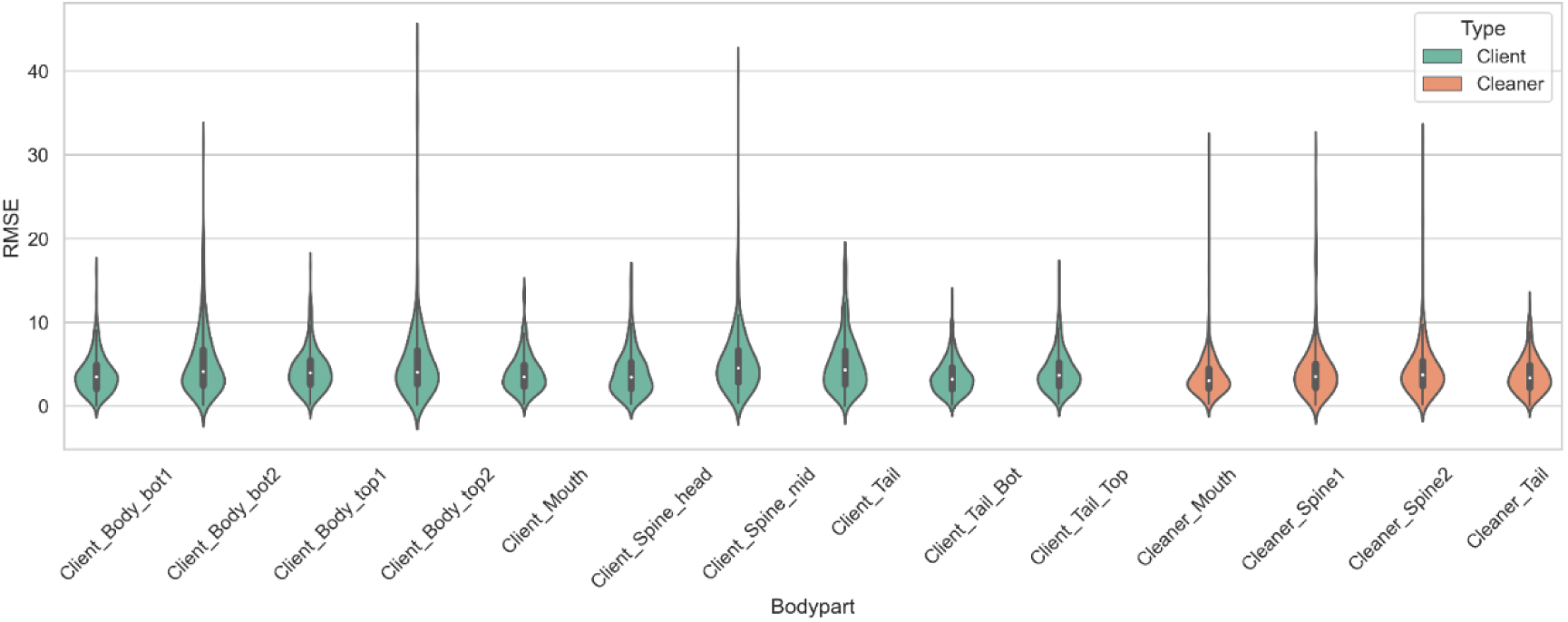
Tracking performance of the Front View model across body parts. The plot displays the mean absolute Euclidean distance (in pixels) for each labelled body part, representing the average deviation between predicted and ground-truth positions. Lower values indicate higher tracking accuracy

#### 3.1.2. Results of TopView setup

The model created for the TopView camera presented a low test error after training (Table S5). The average likelihood for this model was 0.92 and the RMSE values for each body part (as labeled by DeepLabCut) are presented in Figure 5. The low RMSE values and high likelihood indicates that DLC managed to accurately track both fish.

**Figure 5.**
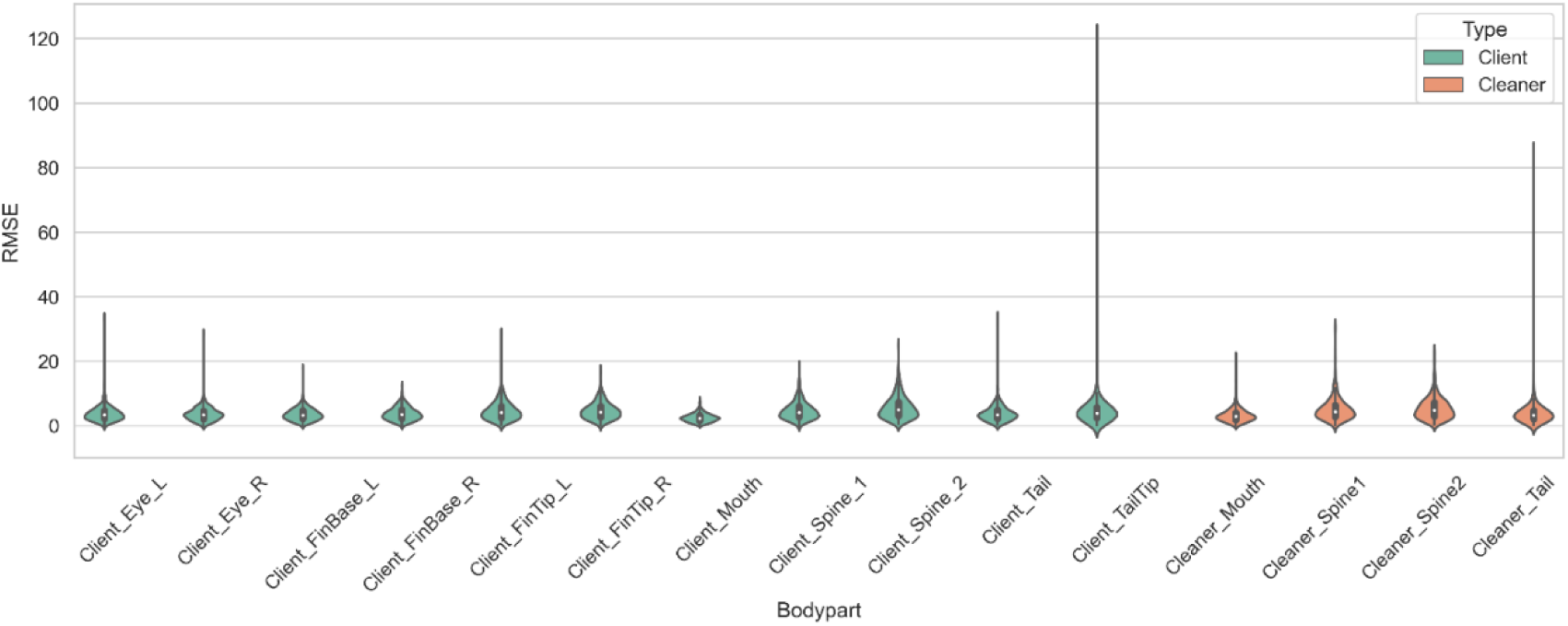
Tracking performance of the Top View model across body parts. The plot shows the mean absolute Euclidean distance (in pixels) for each annotated body part, indicating the average deviation between predicted and true positions. Lower distances reflect higher tracking accuracy for the respective body parts.

### 3.2. Behavioural Classification results

#### 3.2.1. Margin Parameter

No statistically significant differences were found between the models using the parameter *margin* = 0 and *margin* = 30 (p-value > 0.05), suggesting that the choice of margin parameter may not have a substantial impact on model performance in this context (Table 1). Following this, we disregarded the parameter *margin* for the remainder of our results analysis.

**Table 1:**
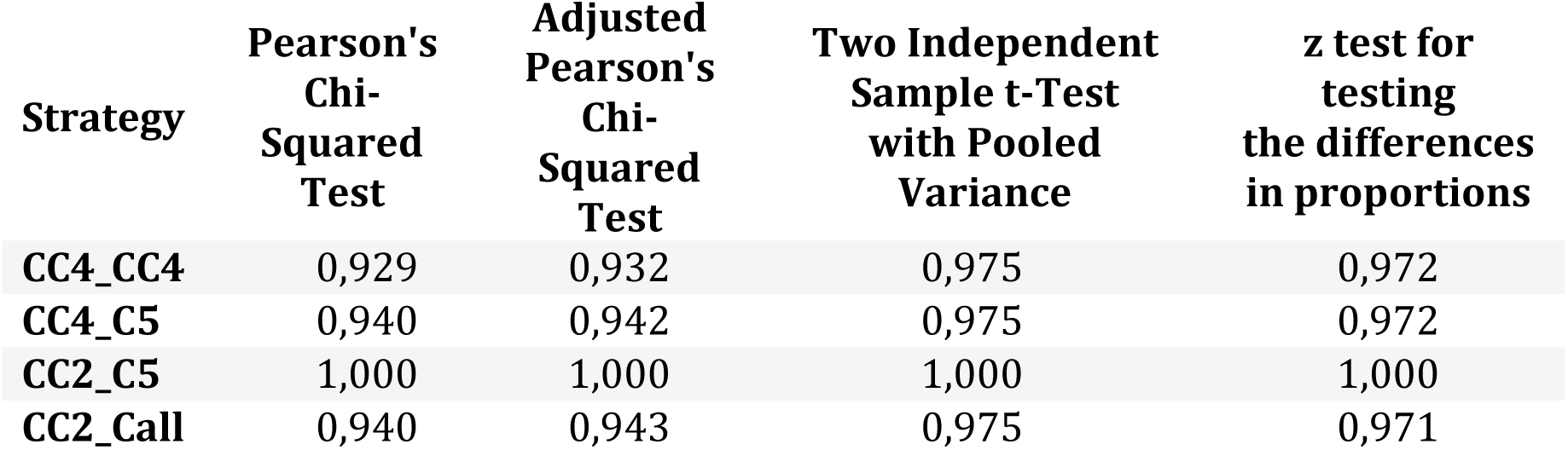
Results of tests comparing proportions of correctly predicted interactions using margin = 0 and margin = 30.

#### 3.2.2. Strategies

All metrics mentioned in sub-section 2.5.2.2 are presented in Table S6, which was used for the analysis of function performance. To better visualize and assess the function’s performance with different parameters, multiple plots were created as shown in Fig. S6. When solely considering the *Min_Frames* change, we observed that all strategies tend to have lower Precision and Specificity but greater F1-score and Recall with lower *Min_Frames* (Figure S6, Table S6).

When focusing on the *Max_Distance* parameter, we observed that Accuracy, F1-Score, and Recall progressively increase when the *Max_Distance* is maximized. However, the opposite occurs for Specificity and Precision, which gradually decrease (Figure S6, Table S6).

We compiled the results of the top-performing model (Strategy 4, *Max_Distance* = 250, *Min_Frames* = 30) into a confusion matrix (Table 2) to give a clearer picture of classification performance. With an accuracy of approximately 87%, recall (true positive rate) of 90%, specificity (true negative rate) 85% and a false-positive rate of 15%, this model demonstrated its robustness to detect real cleaning interactions while minimizing false positives.

**Table 2:** Confusion matrix showing behavioural classification performance of the best-performing model (Strategy 4, *Max_Distance* = 250, *Min_Frames* = 30). The model achieved 90% recall and 85% specificity, with a false-positive rate of approximately 15%. The matrix summarizes predictions across a representative test set of labelled video frames.

|  | *Predicted Interaction* | *Predicted No Interaction* |
| --- | --- | --- |
| *Actual Interaction* | TP = 90% | FN = 10% |
| *Actual No Interaction* | FP = 15% | TN = 85% |

The results from the equation performance showed that, when the *Max_Distance* parameter is increased, the count of Correct Events, Wrong Events, and subsequently the Correct and Wrong Frames increase overall (Table S7). On the other hand, when the *Min_Frames* parameter is increased, the count of Correct Events, Wrong Events, and the predicted Correct and Wrong Frames decreases (Table S7). Additionally, as *Min_Frames* increases, predicted duplicate occurrences become less frequent (Table S7).

## 4. Discussion

We successfully trained two models for pose estimation involving two fish species, *L.* dimidiatus (cleaners) and *A. leucosterno*n (clients), within a controlled laboratory environment and across varied behavioural contexts. To our knowledge, this is the first successful demonstration of multi-animal, full-body pose estimation and tracking in cleaner-client interactions, with consistent model performance across two taxonomically distinct species. Both models were trained on stationary video data where interactions often involved occlusion and complex movement, presenting considerable challenges for automated tracking (A. Mathis et al., 2018; Nath et al., 2019).

Model performance differed between the two camera perspectives. TopView model consistently outperformed the FrontView model in both pixel error and average likelihood. This outcome was expected, as the top-down view presents fewer occlusions of critical body regions than the frontal perspective. The TopView model achieved an average likelihood of 0.96 across all features. Even in scenarios where brief occlusions occurred, such as when fish swam beneath the overflow system (See supp. figure S7), the model maintained stable tracking. The FrontView model, by contrast, exhibited a higher average error (8.66 pixels), though this remains small relative to the 1920×1080 resolution of the videos. The reduced average likelihood for the FrontView model (0.92) is primarily due to natural occlusions—e.g., one fish swimming in front of the other or turning away from the camera (Supp. Figure S8A– B) a common issue in three-dimensional tracking setups (Nath et al., 2019). Our models achieved pixel errors similar to other studies using DLC on fish (Ravan et al., 2023), validating the robustness of our training process and reinforcing DLC’s applicability in complex aquatic systems. Moreover, when applied to entirely novel videos not included in training, the models continued to perform well, as verified through visual inspection and the provided example videos (Supp. Video X). These results indicate that our approach generalizes effectively to new datasets.

Although our pose estimation pipeline was successful, behavioural classification proved more challenging. Unlike prior studies focused on single animals in 2D settings (e.g., Bohnslav et al., 2021)), our work involved multiple interacting individuals in a 3D aquatic environment, limiting direct comparability. Nevertheless, our classification function significantly reduced the workload of manual video analysis. Using Strategy 4, with a *Max_Distance* of 250 and *Min_Frames* of 30, we reduced the number of frames requiring human validation from around 500,000 (about 4 hours and 40 minutes of footage) to between 104,000 and 126,000 frames (roughly 57 to 70 minutes). This represents a 75% reduction in total analysis time, with only around 10% of interactions missed, most of which could be recovered by slightly expanding the window around predicted events.

This semi-automated approach offers a practical and scalable solution for researchers working with large video datasets, particularly in systems characterized by rapid and frequent interactions, such as cleaning mutualisms. Manually coding such behaviour is highly time-intensive: a one-hour video may take 45 minutes to annotate if interactions are sparse and the observer has the video sped up, or several hours if densely populated. Our reduction in processing time marks a substantial advance toward scalable behavioural analysis with limited human resources.

The results support the feasibility and value of deep-learning-based posture estimation in multi-species aquatic systems under naturalistic behavioural conditions. The strong model performance both during training and when generalizing to unseen data demonstrates the effectiveness of our dataset design and frame selection process. That said, the model-building process remains time-consuming and could hinder wider adoption. Future efforts should aim to enhance generalizability by expanding training datasets across varied environmental contexts, including different tank layouts or additional species. Behavioural classification could also be improved by incorporating supplementary information such as movement trajectories, acceleration data, or proximity metrics. Collaborative efforts to share annotated datasets across research groups would further support this goal.

In conclusion, we demonstrate that multi-species posture estimation and tracking of cleaning interactions is not only feasible but highly effective when grounded in robust training data and realistic video inputs. Moreover, our pipeline offers a substantial improvement over manual scoring for the detection of cleaning interactions, laying the groundwork for more scalable, reproducible, and quantitative studies of social interactions.

## Acknowledgements

We thank all members of the Behavioural Ecology and Evolution Group and Eve Otjacques for stimulating discussions and comments on this manuscript.

---

**Figure S1:**
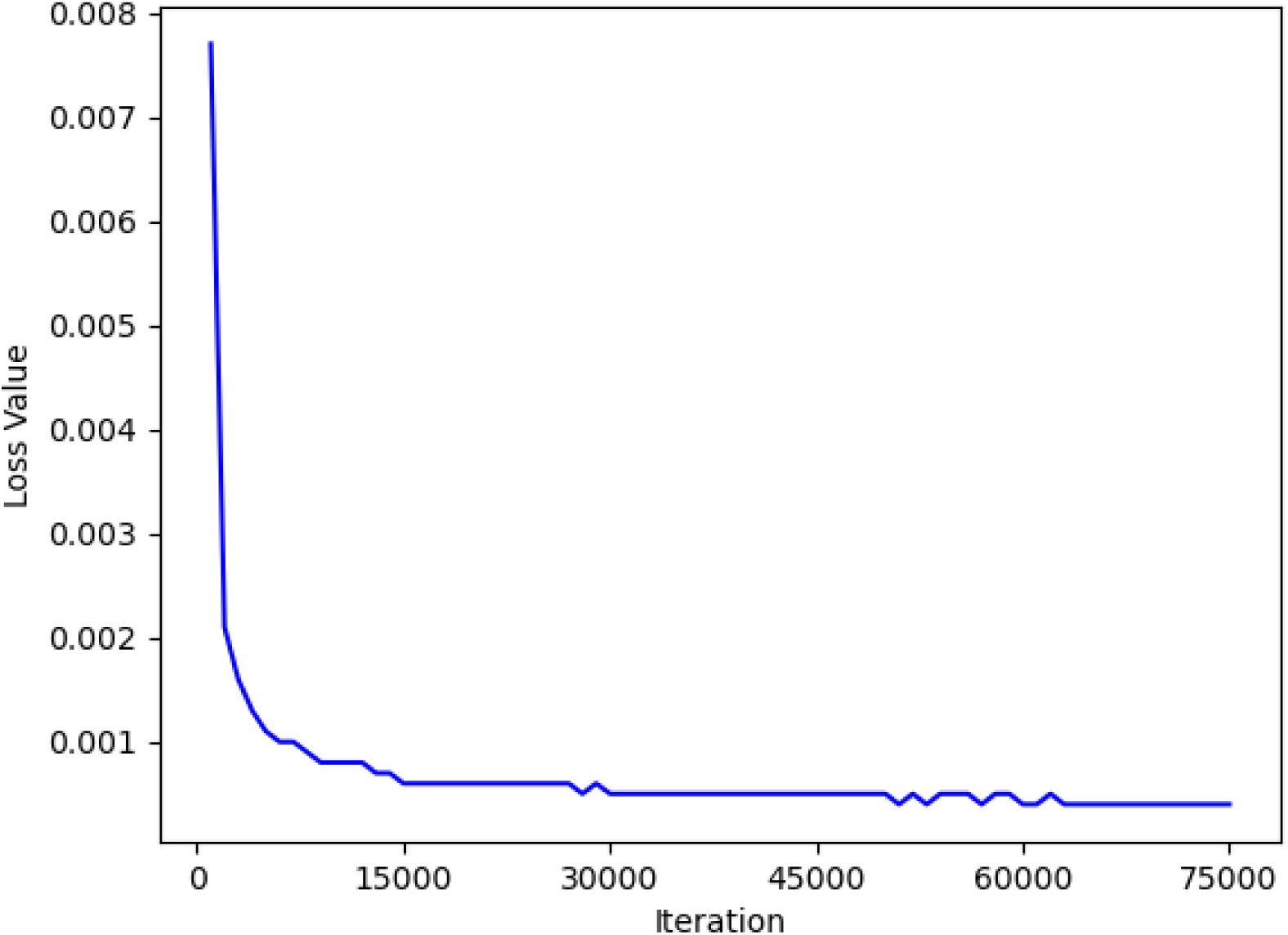
Training loss curve of the DeepLabCut model for the front-view camera. The plot shows the reduction in loss value over training iterations, indicating the pr ogressive improvement in model performance during the training process.

**Figure S2.**
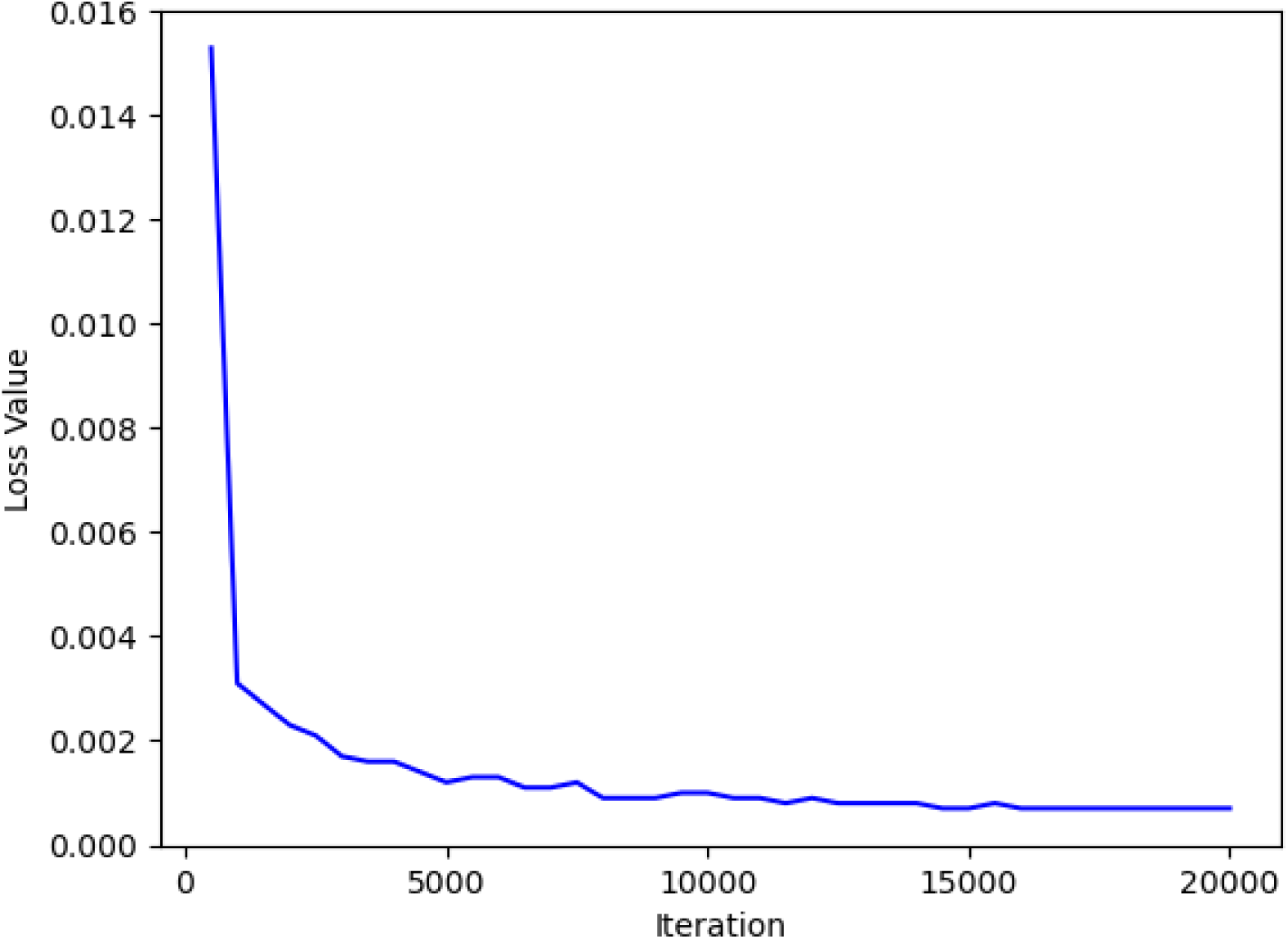
Training loss curve of the DeepLabCut model for the top-view camera. The graph illustrates the decline in loss value over training iterations, indicating the model’s convergence and improvement in tracking accuracy throughout the training process.

**Figure S3:**
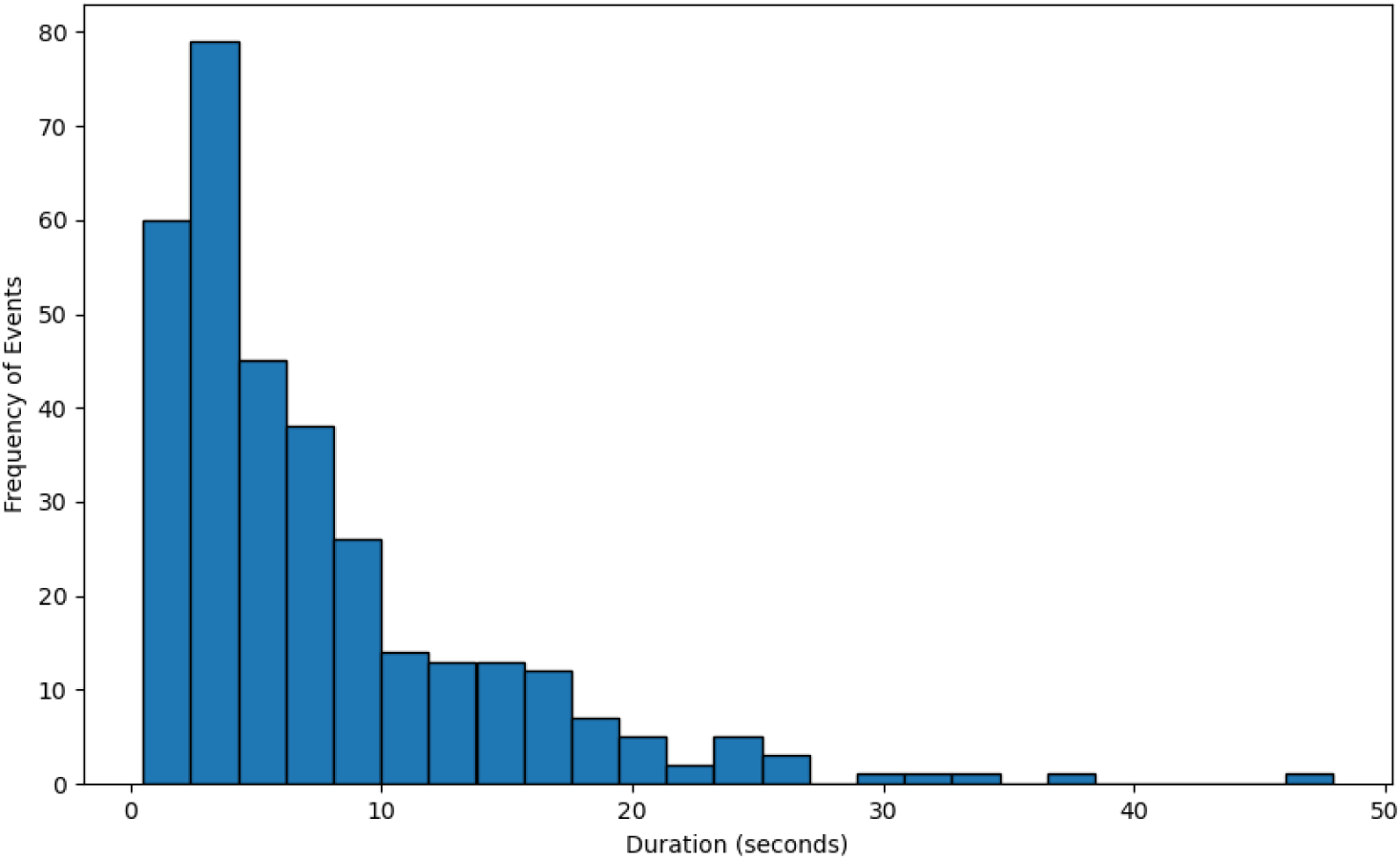
Frequency distribution of interaction duration (seconds) of all videos

**Figure S4:**
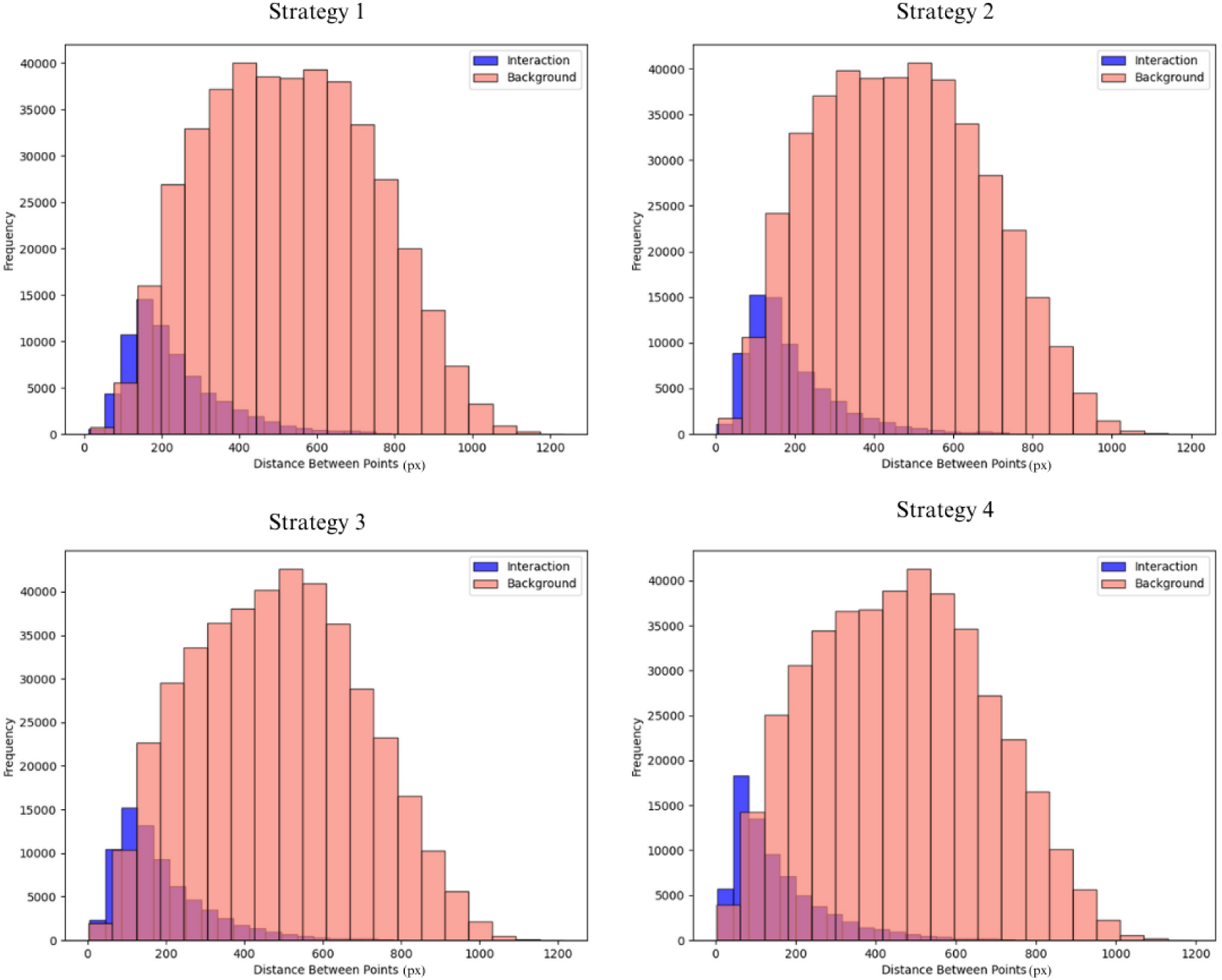
Comparative Distribution of Distance Measurements (px) across the four Strategies used, Interaction vs Background

**Figure S5.**
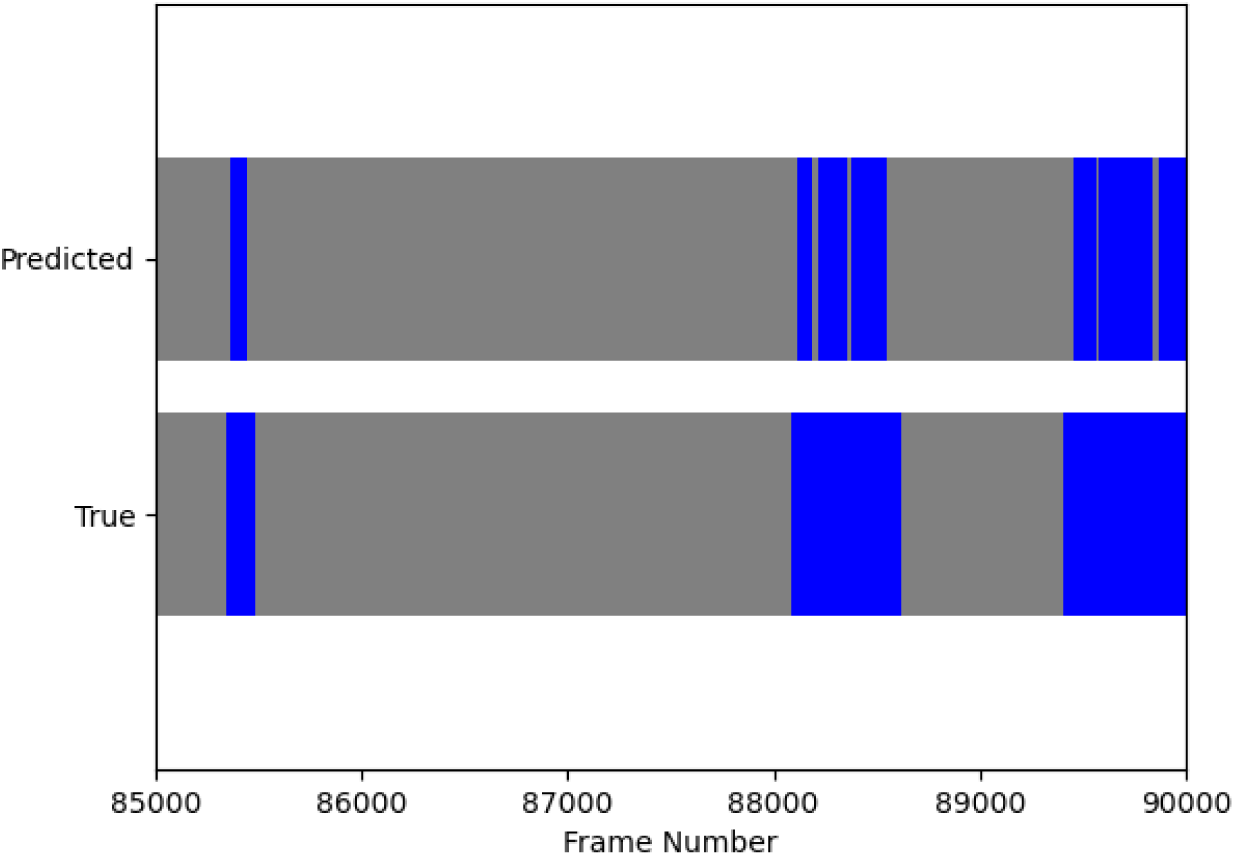
Example of duplicate predicted interaction event (blue). The behavioural classification model identified seven interaction events whereas only three actual events occurred in the video. This resulted in four false-positive duplicates, where a single true interaction was mistakenly segmented into multiple predicted events.

**Figure S6.**
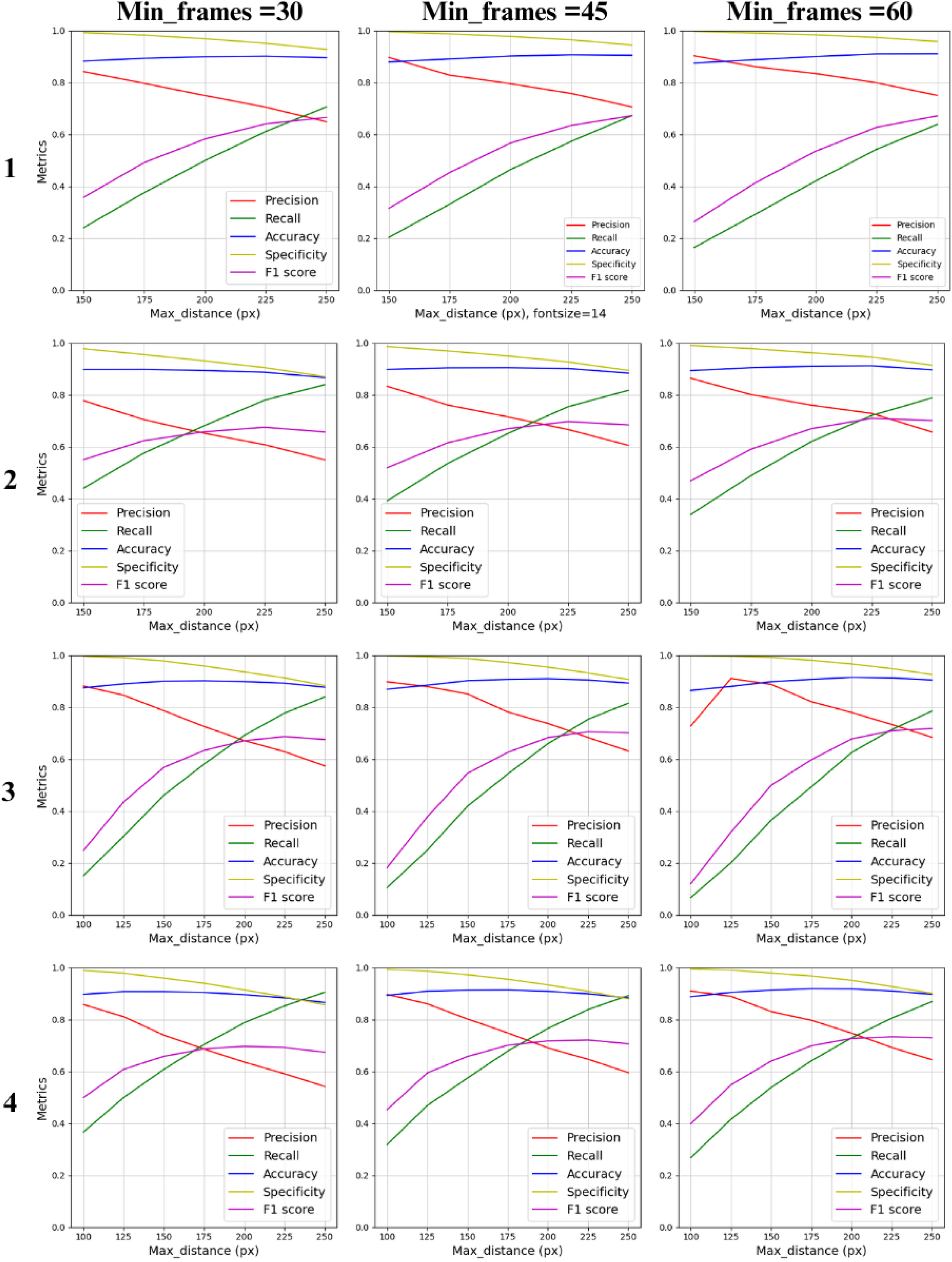
Performance comparison of four classification strategies across function parameters. Each panel corresponds to a different Min_Frames threshold (30, 45, and 60 frames, from left to right). Within each panel, the classification performance of strategies 1 through 4 is compared across varying Max_Distance values. This visualization highlights how parameter combinations influence the detection of cleaning interactions

**Figure S7:**
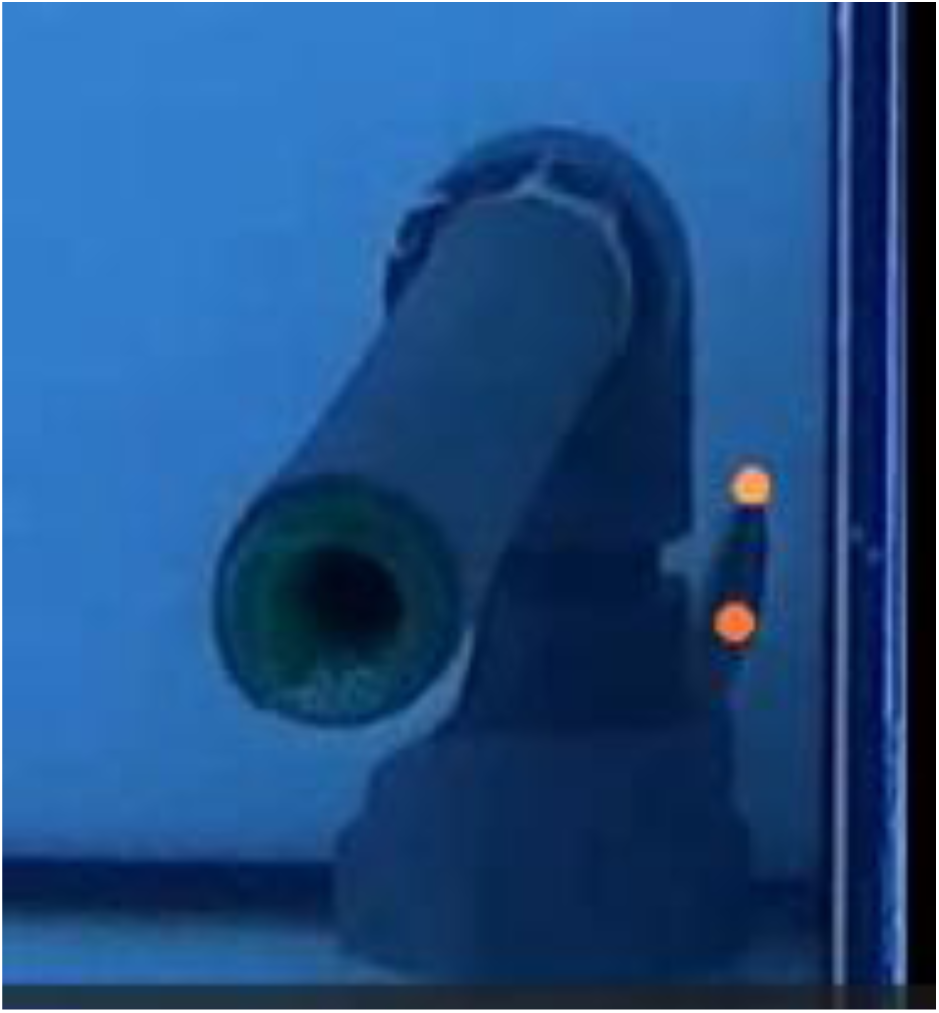
*Labroides dimidiatus* hidden under the overflow system, making it difficult to track

**Figure S8:**
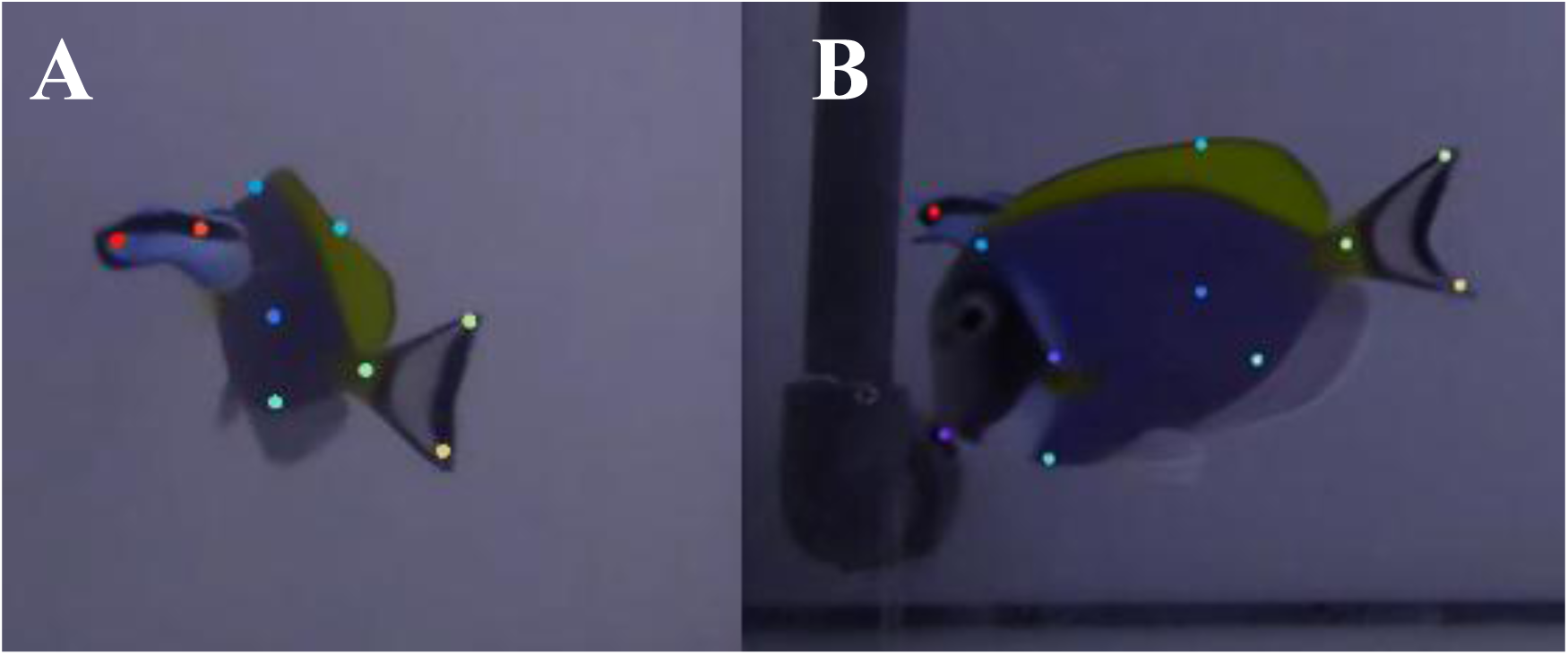
FrontView untraceable examples. A-Both fish swimming away from the camera. B-Moments before the *Labroides dimidiatus* being occluded for brief moments.

**Table S1:** Fish IDs and corresponding Metadata.

| Fish ID | Duration (min) | Camera View |
| --- | --- | --- |
| LD03 | 62 | Top |
| LD04 | 52 | Top |
| LD13 | 54 | Top |
| LD14 | 56 | Top |
| LD23 | 63 | Top |
| LD24 | 48 | Top |
| LD03 | 50 | Front |
| LD04 | 63 | Front |
| LD13 | 62 | Front |
| LD14 | 46 | Front |
| LD23 | 50 | Front |
| LD24 | 56 | Front |

**Table S2:** Interaction Frequency Distribution by Time Intervals.

| Duration range<br>(seconds) | Occurrences | Cumulative |
| --- | --- | --- |
| [0, 1] | 5 | 5 |
| ]1, 1.5] | 16 | 21 |
| ]1.5, 2] | 43 | 64 |
| ]2, ∞ [ | 284 | 327 |

**Table S3:**
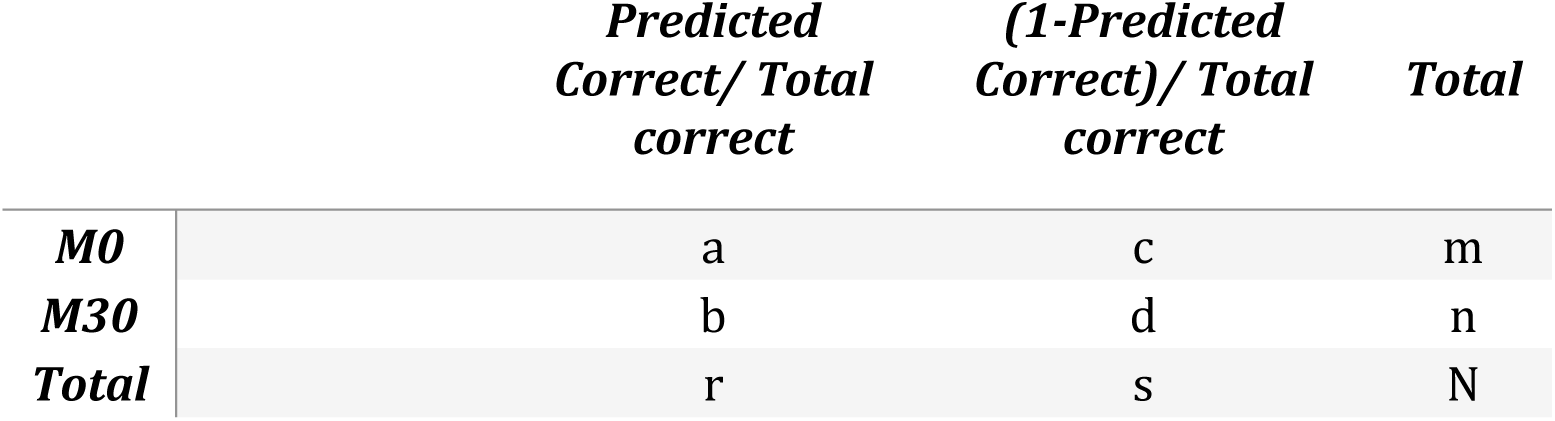
The observed frequencies in a 2x2 Table.

**Table S4.** Training and test error summary at iteration 75,000 for the Front View model. The table compares the mean tracking error (in pixels) between the training and test datasets, reported both with and without applying a p-cutoff threshold of 0.75.

|  | Train | Test |
| --- | --- | --- |
| Error (px) | 4.02 | 8.66 |
| Error with p-cutoff (px) | 4.02 | 8.32 |

**Table S5.**
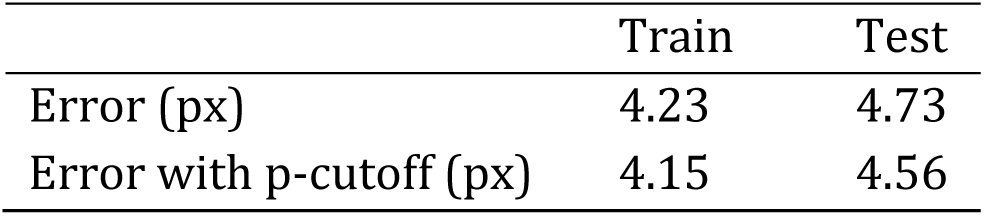
Training and test error summary at iteration 20,000 for the Top View model. The table compares the mean tracking error (in pixels) between the training and test datasets, reported both with and without applying a p-cutoff threshold of 0.75.

**Table S6.**
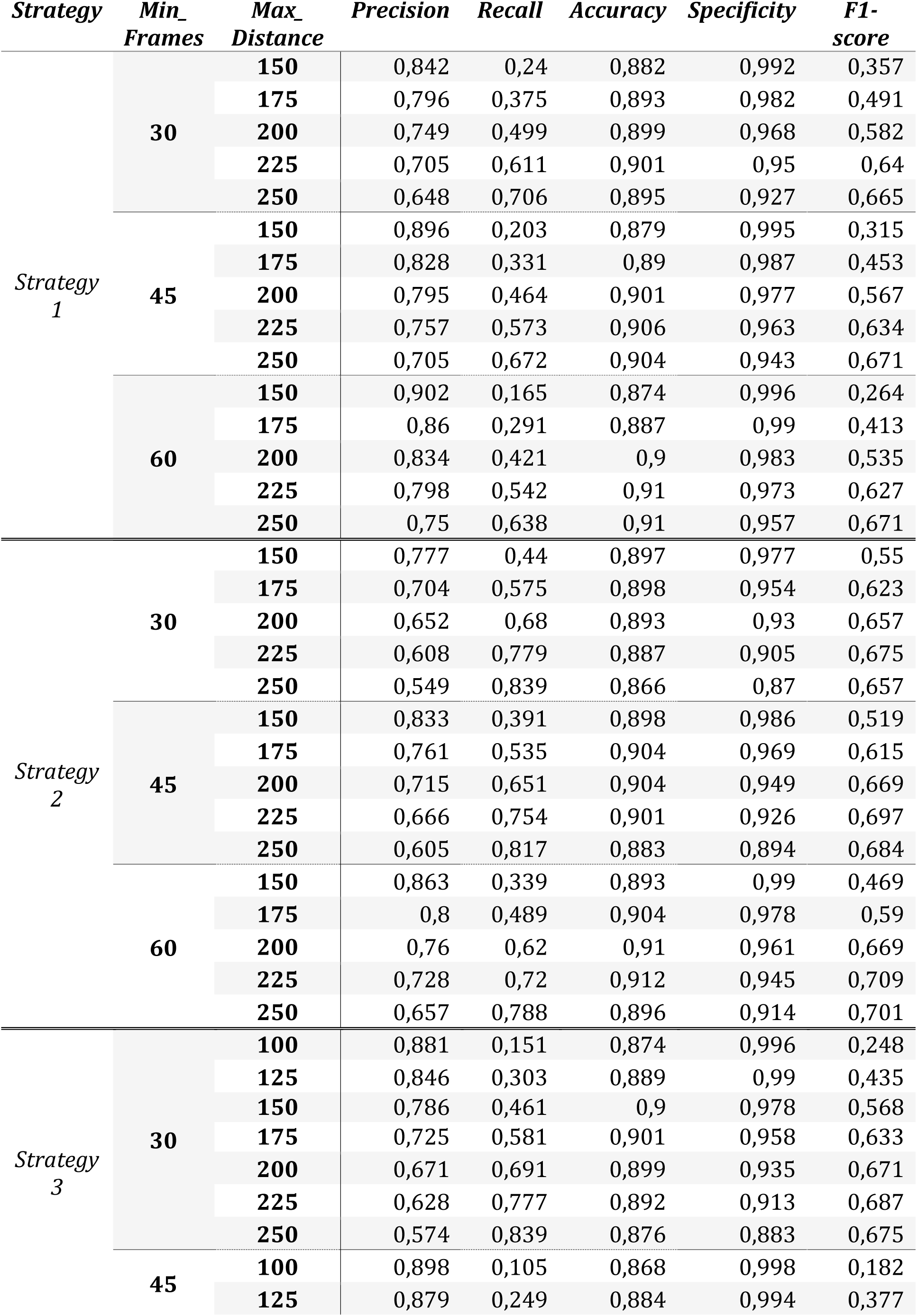
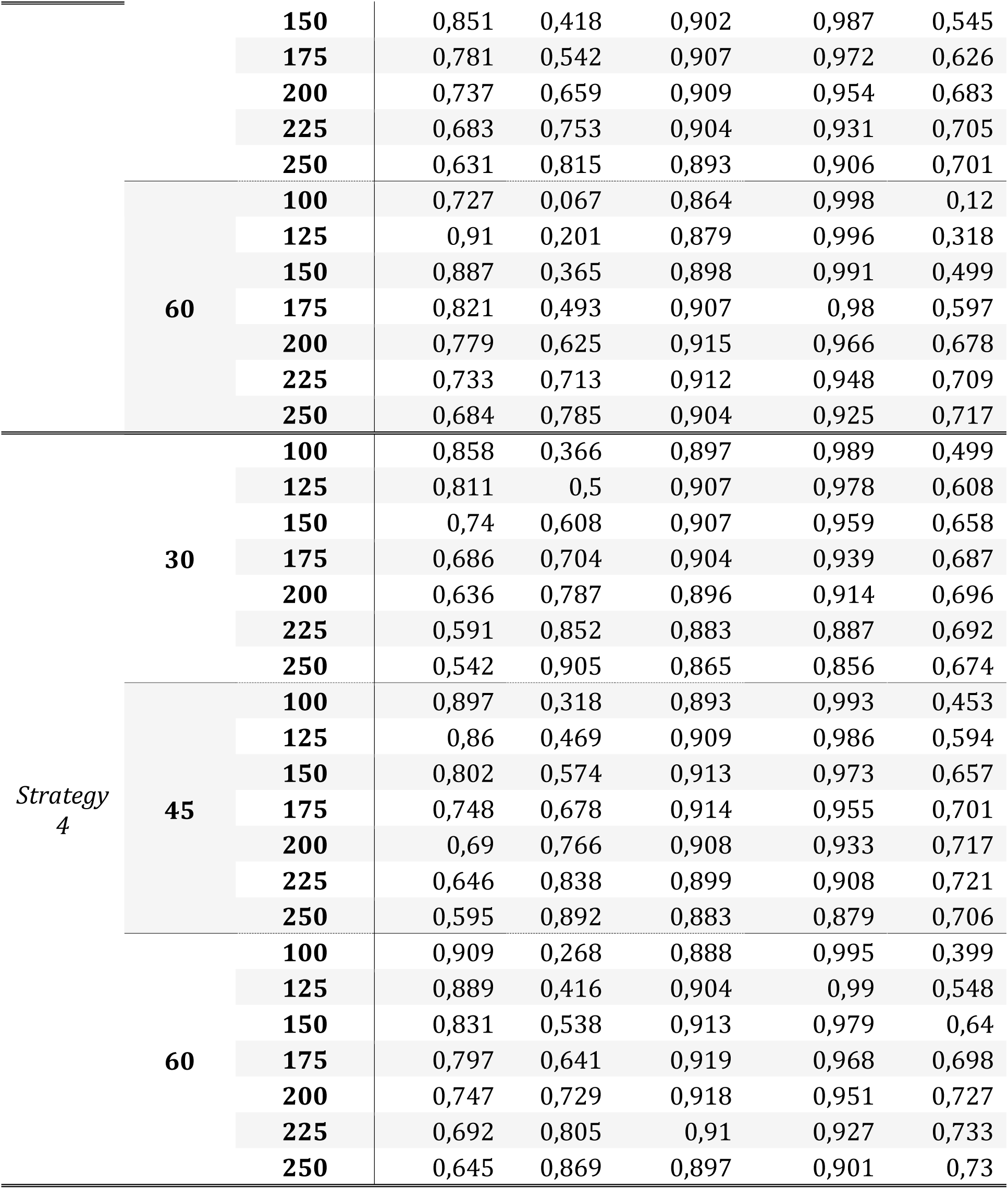
Function evaluation Metrics.

**Table S7.**
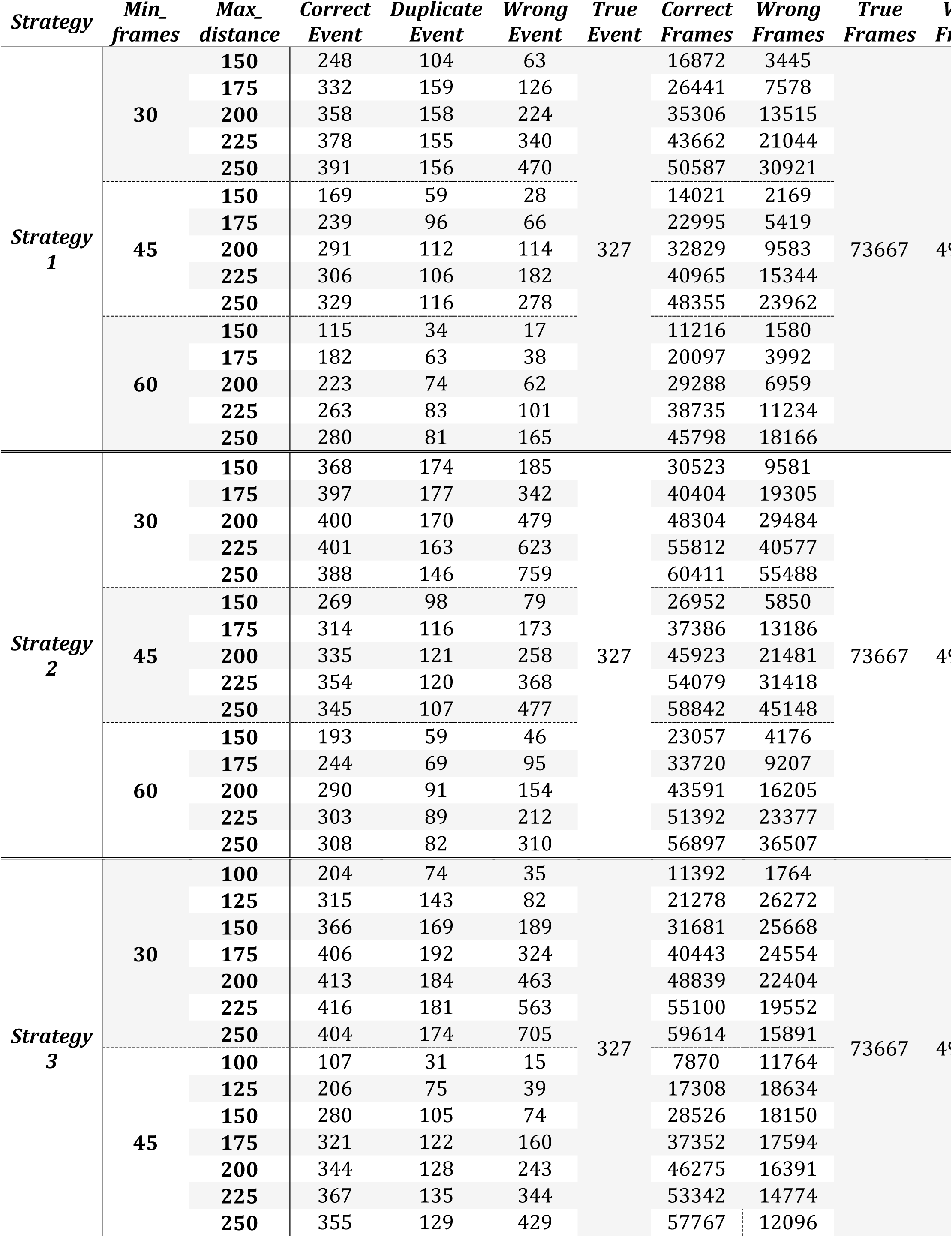
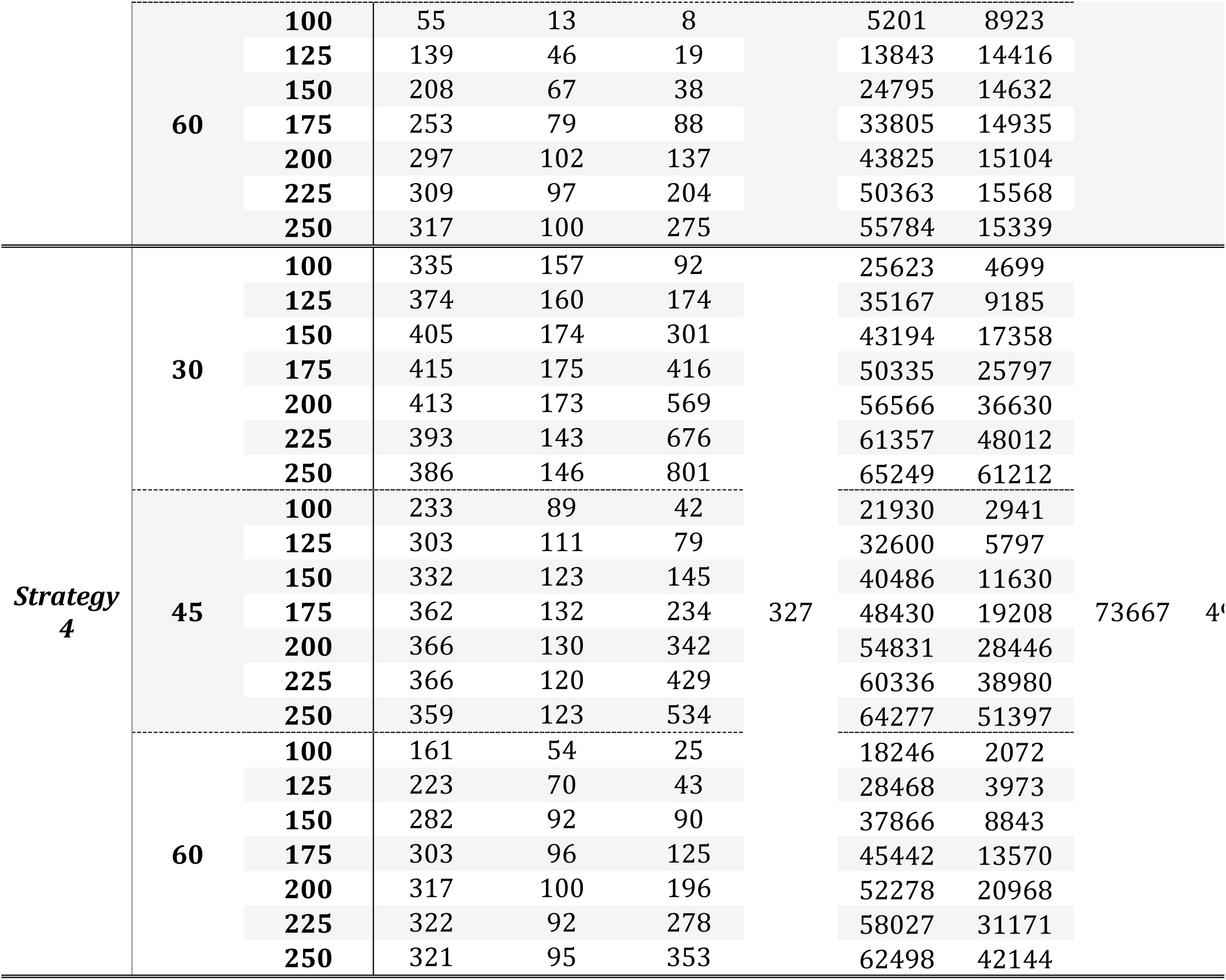
Function evaluation total predictions.

